# Lymphoid-tissue-on-chip recapitulates human antibody responses in vitro

**DOI:** 10.1101/2025.01.14.632762

**Authors:** Claudia Teufel, Anna-Sophie Schlemmer, Andrea W. Eiken, Zachary W. Wagoner, Dennis Vöhringer, Lena Christ, Alex Dulovic, Patrick Marsall, Madeleine Fandrich, Eduardo J.S. Brás, Julia Marzi, Friederike Bärhold, Julia Philipp, Sven Becker, Lisa E Wagar, Peter Loskill

## Abstract

In the past decades, vaccine development has made great strides. Nevertheless, more often than not, vaccine candidates fail in advanced stages of development and clinical trials. A key reason is the poor predictive value of non-clinical in vivo and in vitro models, due to either species-specific differences in the immune response or insufficient reflection of physiological vaccine mechanisms. Reliable modeling of human adaptive immune responses is a prerequisite to understand processes leading to vaccine-induced protective immunization and to drive informed decisions in vaccine development pipelines. Here, we present a centrifugal microfluidics based organ-on-chip approach to generate an organotypic high density lymphoid tissue on-chip. The model enables long-term culture of lymphoid tissue and raised antigen-specific antibody responses against influenza vaccines even after four weeks on-chip. Antibody response of different magnitude and quality could be induced both by direct antigen exposure as well as by recruitment of antigen-presenting cells from the periphery. The model represents an attractive approach to evaluate the impact of the mode of antigen delivery on adaptive immune responses. Beyond applications in vaccine development, the lymphoid-tissue-on-chip provides a platform to study cellular interactions during homeostasis, immune responses and long-term impact of immunomodulators over several weeks.

## INTRODUCTION

Vaccinations are the best strategy to control endemic and epidemic infectious diseases and to reduce the cumulative impact of infection-associated morbidities. Through the inclusion of pathogen-specific target materials, vaccines stimulate an adaptive immune response resulting in the production of antigen-specific antibodies and T cells. The T cell-dependent antibody response is a multi-step process that results in the development of B and T cell memory and long-lasting antibody responses (*1*). This process involves both antigen-presenting cells from peripheral organs, in particular dendritic cells, as well as stromal cells and immune cells located in secondary lymphoid organs such as lymph nodes and tonsils (*2–4*). Upon vaccination, peripheral and lymphoid tissue-resident dendritic cells become activated by adjuvant-mediated signals through pattern recognition receptors, take up the vaccine antigens, and peripheral dendritic cells migrate to lymphoid organs (*5*). In the lymphoid tissues, the interplay of activated dendritic cells, antigen-specific T follicular helper cells, other antigen-specific T cells, innate immune cells and stromal cells promotes an early extrafollicular B cell response and the formation of germinal centers (*4*, *6–8*). Direct cell-cell contacts and released factors support B cell survival, proliferation, class switching and antibody affinity maturation as well as B cell differentiation into germinal center B cells, memory B cells, short-lived plasmablasts and long-lived plasma cells (*3*). Protective antibodies generated during germinal center reaction are highly specific, neutralize pathogens and guide immune cell effector functions.

Most of the knowledge on antibody responses originates from in vivo animal studies, predominantly used for non-clinical vaccine development. However, many vaccines against e.g. *Mycobacterium tuberculosis* (*9*) or malaria (*10*) fail during clinical trials due to poor immunological responses or efficacy within their target population. A retrospective cohort study on the probability of success of vaccines developed against emerging and reemerged viral infectious diseases concluded that only one in ten vaccines progresses from clinical phase II studies to U.S. Food and Drug Administration approval within ten years (*11*). This probability is even reduced to one in 30 vaccines if influenza vaccines produced in well-established platforms are excluded (*11*). Efficacy is also a concern among routinely used vaccines due to its variable protection in different demographic populations, as exemplified in different hepatitis B vaccine response rates in young versus adult individuals and further confounding factors within the adult population (*12*). The poor predictive value of animal vaccine testing originates from described ‘nonmodifiable discrepancies’ between humans and experimental animals, such as inherent differences in the immune system, immunological history, susceptibility to pathogens and pathophysiology, which cannot be overcome by new strategies to increase comparability of data from experimental animals to the human system (*13*). Therefore, there is an urgent need for human in vitro models which emulate complex immunological processes such as vaccine-induced antibody response.

In recent years, the development of human 2D and 3D lymph node models that reflect functional aspects of central immune responses has undergone significant progress (*14*). Models range from simple immune cell co-culture systems with or without hydrogel, to static and microfluidic culture of lymph node tissue sections and bioreactors and microfluidic platforms (*14*, *15*). Many of these models recreate individual aspects of immune responses and have proven to be useful to elucidate immune cell migration behavior in signal gradients or the core aspects of germinal center reaction (*15*). However, most of these models rely on blood-derived immune cells with or without artificial stromal cell lines and therefore lack the complexity of stromal and immune cell heterogeneity necessary to reproduce T cell-dependent antibody responses. Models with more cellular complexity and advanced spatial microanatomical organization such as lymphoid tissue sections are limited in application due to their short viability of up to seven days (*16–18*). Furthermore, most models ignore the functional impact of cell density on immune cell function (*19*). On the cellular level, the most advanced lymphoid tissue model has been developed by Wagar et. al (*20*) and consists of tonsil aggregates composed of stromal and immune cells derived from human tonsil biopsies. Tonsil aggregates displayed lymph node-like spatial cell distribution, mounted antigen-specific antibody responses against different types of vaccines in a T cell-dependent manner and recapitulated many characteristics of germinal center B cell differentiation and antibody maturation, including somatic hypermutation and class switching. Drawbacks are limited culture longevity with a donor-dependent decline in B cells after two to three weeks in culture and the lack of physiological delivery of nutrients, compounds and particularly antigens, e.g., via peripheral antigen-presenting cells. In recent years, organ-on-chip technology has emerged as a promising tool to generate microphysiological tissue models by modifying the culture environment to meet tissue-specific cell requirements and by enabling a vasculature-like perfusion, thereby supporting longevity of various tissue models, enhancing tissue function and allowing for perfusion with different types of immune cells (*21–23*).

Here, we merged organ-on-chip technology with the tonsil aggregate approach to generate a microfluidic human lymphoid-tissue-on-chip (LToC) model, which enables high-density culture of lymphoid tissue-derived immune and stromal cells, increased culture longevity and different modes of antigen delivery (Fig. 1). Core features of the model include a specifically designed centrifugal microfluidic chip for high-density lymphoid tissue generation, vasculature-like perfusion for up to four weeks of on-chip culture and robust assays for vaccination studies highlighting functional antibody-response upon stimulation directly with antigens or indirectly via autologous antigen-presenting monocyte-derived dendritic cells.

**Fig. 1:**
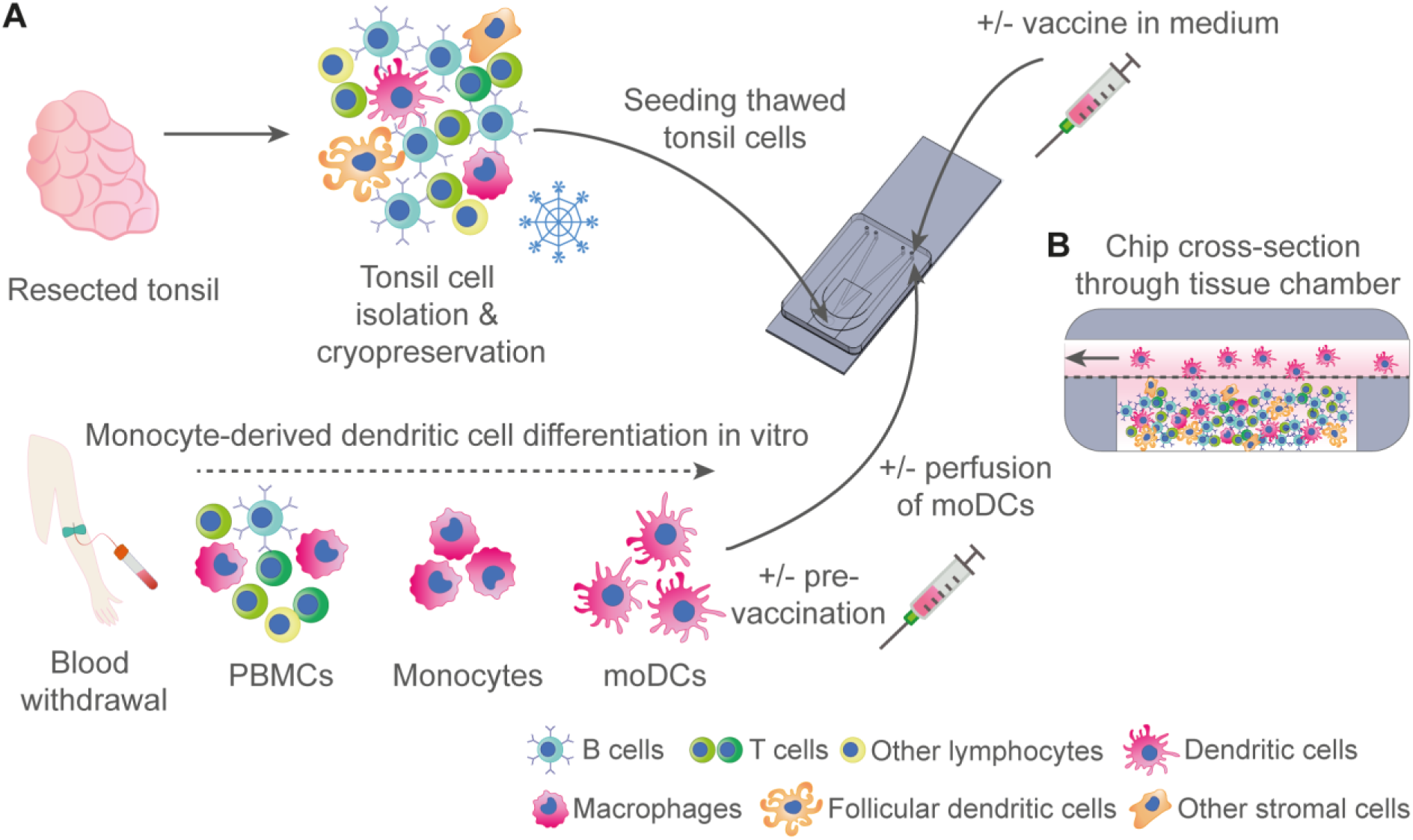
Concept of the lymphoid-tissue-on-chip (LToC) model with adjustable mode of antigen delivery. **A** Schematic of cell isolation, monocyte-derived dendritic cell (moDC) differentiation and cell component integration into the LToC platform. **B** Schematic of LToC chamber filled with tonsil-derived cells and recruitment of perfused moDCs into the tissue-chamber.

## RESULTS

### Centrifugal microfluidic lymphoid tissue chip enables hydrogel-free, high-density culture of tonsil-derived lymphoid tissue

To enable seeding and culture of human lymphoid tissue-derived cells at high densities, we developed a fit-for-purpose lymphoid tissue (LT) microfluidic chip. In the LT chip, cells can be compacted via centrifugation into a tissue chamber (3 mm diameter, 0.3 mm height), which is separated from an overlying medium channel (0.2 mm height) by a porous polyethylene terephthalate (PET) membrane with 5 μm pores (Fig. 2A). The chosen pore size retains all cells within a confined area upon centrifugation, but at the same time allows for active transmigration of immune cells (*22*, *24*). Human tonsil biopsies, which are easily accessible and contain all relevant stromal and immune cell subsets involved in T cell-dependent antibody responses (*25*), were used as a source for lymphoid tissue-derived cells and dissociated using a previously established enzyme-free dissociation protocol (*20*) (Fig. 2B). To generate lymphoid-tissue-on-chip (LToC), cryopreserved tonsil cells were thawed, injected into the tissue seeding channel of LT chips, and condensed into the tissue chamber by centrifugation (Fig. 2C). Then the tissue seeding channel was sealed with Dextran CD hydrogel. Upon unidirectional medium perfusion through the medium channel (40 μL/h) with a syringe pump, tonsil cells populated the tissue chamber in multiple layers and reaggregated into clusters of more condensed regions containing B and T cells (Fig. 2D), similar to tonsil aggregates in static plate cultures (*20*). All in all, the LT chip provides an environment in which lymphoid tissue can be generated from tonsil-derived cells at high density, with continuous medium flow that does not interfere with their capacity for self-aggregation.

**Fig. 2:**
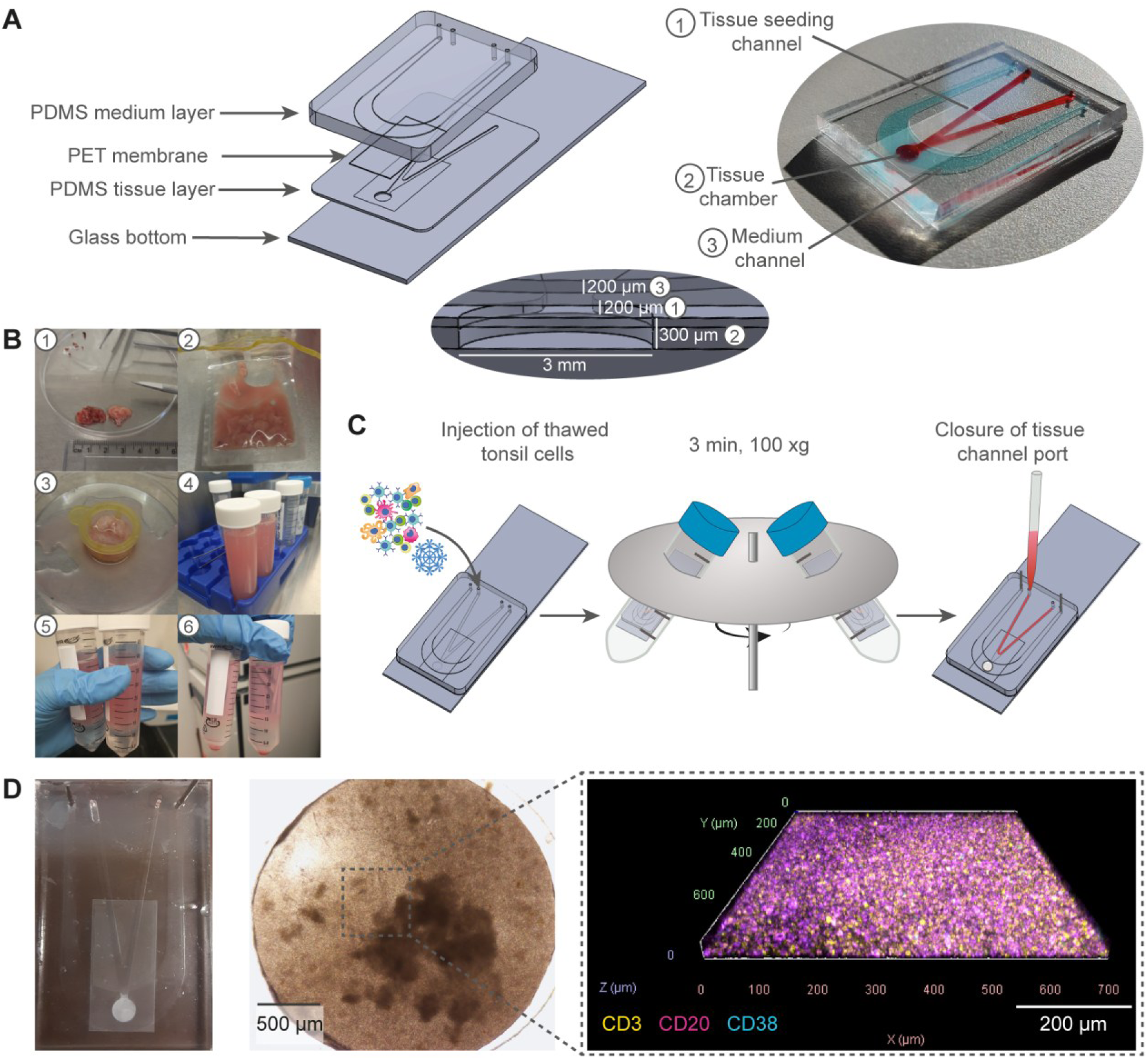
Lymphoid-Tissue-on-Chip (LToC) design and cell injection. **A** Schematic depiction of the four chip layers incorporating polydimethylsiloxane (PDMS) tissue and medium layer, a porous polyethylene terephthalate (PET) membrane and a glass bottom (microscope slide) (left). Dimensions of tissue chamber and media layers (middle). Water color-filled, assembled LToC with coverslip glass bottom (right). **B** Images depicting the tonsil isolation process: 1) Washed tonsil parts, 2) chopped tonsil fragments after first processing, 3) tonsil fragments after repeated smashing over cell strainer, 4) tonsil suspension before centrifugation, 5) + 6) tonsil cell suspension before and after gradient centrifugation. **C** Schematic of centrifugal cell injection into the LToC. **D** Exemplary overview image of LToC after nine days under perfusion (left), bright-field microscope close-up image of tissue chamber (middle) and immunofluorescence 3D image of CD20^+^ B cells, CD3^+^ T cells and CD38^+^ cells.

### LToC remains viable and maintains immune cell subset stability over four weeks of perfused culture

The lifespan of statically cultured tonsil aggregates is limited to about three weeks, but the number of recovered living cells already drastically decreases between two and three weeks of culture for some donors (*20*). To determine the longevity of the perfused LToC, we monitored the viability and immune cell composition of unstimulated LToC over the course of four weeks. Effluent quantification of lactate dehydrogenase (LDH), a surrogate cell death marker, showed largely stable LDH levels over time, indicating no major cell death events occurred throughout the culture period (Fig. 3A). Staining with Calcein AM viability dye and propidium iodide cell death counterstaining on weekly sacrificed chips showed that large numbers of cells were viable over four weeks (Fig. 3B). To assess cell viability and to perform immune cell subset analysis, we retrieved cells from LT chips one day after seeding and subsequently on a weekly basis, and subjected the retrieved cells to flow cytometric analysis. The absolute number of recovered cells (based on flow cytometry data) was stable over the entire culture period (Fig. 3C). Viability of cells retrieved from LToC gradually decreased over the first three weeks of culture and stabilized at a viability of about 50% of the initially viable cells after three weeks. The percentage of hematopoietic cells (CD45^+^) was maintained around 98-99% and the overall proportions of B cells (CD19^+^), T cells (CD3^+^), and non-B, non-T cells (e.g. NK cells, dendritic cells) were comparable across all time points, with only a small decrease in the frequency of T cells over time. Next, we wanted to determine whether the cells within the LToC preserve functional capacity throughout culture. We therefore collected effluents every second day and applied four exploratory multiplex panels covering a total of 68 lymphoid tissue-associated cytokines, general immune-related cytokines, complement components, extracellular matrix proteins, soluble Fas and soluble cytokine receptor isoforms in LToC effluent collected at five different timepoints to measure secretion and shedding of these analytes by LToC over time (Fig. 3D, Fig. S1). 29/68 of the measured analytes were below analyte-specific lower limit of quantitation, 3/68 analytes (aggrecan, chemerin, IL-15) were detectable at equivalent concentrations in LToC effluents and medium control and BAFF, which was supplemented in the medium, was above upper limit of quantitation. The concentration of 35/68 analytes in effluent from LToC exceeded medium level concentrations. These included lymphoid tissue-associated cytokines like CXCL13, CCL19, IL-17 and interferon (IFN)-α, immunomodulatory soluble B cell maturation antigen (BCMA) (*26*) and Collagen IVα1. Most measured analytes were either stable or decreased in parallel to the overall viability of the LToC. The majority of stably expressed analytes comprised chemokines and interleukins, suggesting continuous signaling among tonsil-derived cells. Only some activation- or stress-related analytes like IL-1ra, IL-6R-α, IL-8 and IL-10 were exclusively detected on day 2, which might reflect cellular stress post-thawing. These findings indicate that the heterogenous population of tonsil cells in the LToC retained the ability to produce lymphoid tissue-relevant cytokines and other secreted factors for four weeks. Taken together, this data demonstrates that the LToC has extended viability compared to other approaches to culture lymphoid tissue, shows a robust ratio of B and T cells throughout culture and shows continuous production of lymphoid tissue-associated factors.

**Fig. 3:**
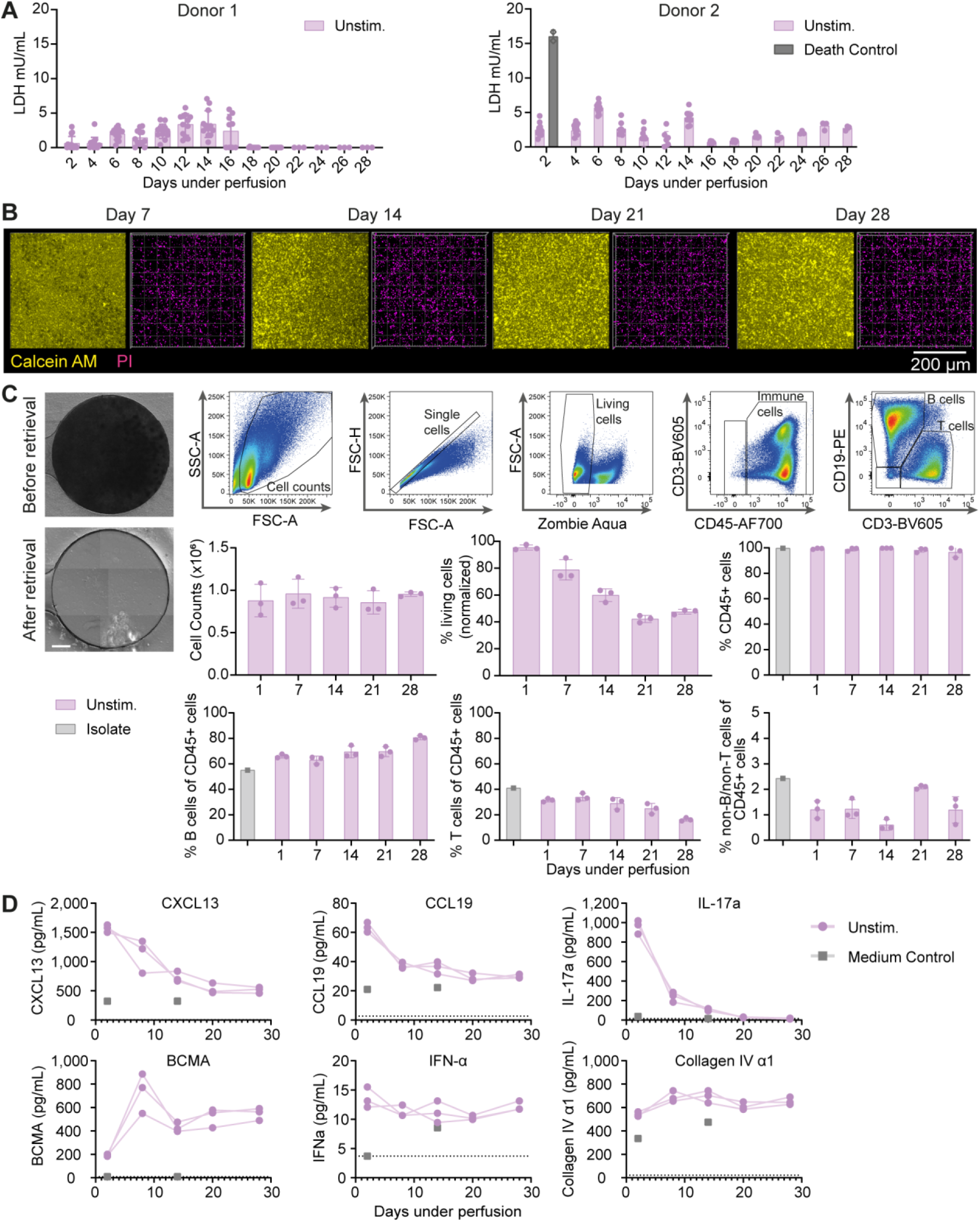
LToC viability, cell subset distribution and protein secretion during four weeks of on-chip culture. **A** Lactatdehydrogenase (LDH) measurements as cell death surrogate marker in effluents collected every other day from unstimulated (unstim.) LToCs or cell death control (1 % Triton X-100). n = 3-15 chips (unstim.), n = 2 chips (death control) (two donors). **B** Live-dead-staining of unstimulated LToC terminated at 7, 14, 21 and 28 days under perfusion. Exemplary 3D top view images of regions within the LToC tissue chamber stained with Calein AM (living cells) and propidium iodide (PI) (dead cells). **C** Cell retrieval from unstimulated (unstim.) LToC for longitudinal flow cytometric analysis of cell subset distribution. Images show one exemplary LToC chamber before and after cell retrieval (left, scale bar 500 μm) and the flow cytometry analysis gating strategy (right). Cell counts were estimated from flow cytometry cell counts. Cell viability was normalized to initial viability of tonsil isolates. Ratios of immune cells (CD45^+^ cells), B cells (CD19+), T cells (CD3+) and non-B/non-T immune cells in LToC after 1, 7, 14, 21 and 28 days under perfusion and in tonsil isolate before chip loading (isolate). n = 3 (unstim.) and n =1 (medium control). **D** Multiplex cytokine analysis of LToC effluents (unstim.) and medium control. Effluents were collected every other day and effluents from day 2, 8, 14 and 28 were measured for secretion of chemokines (CXCL13, CCL19) cytokines (IL-17, IFN-α), B cell maturation antigen (BCMA) and Collagen IV α1). Each continuous line depicts effluents from one LToC. Dashed lines indicate lower limit of quantitation of respective analyte. Medium controls were measured from day 2 and day 14.

### B cells differentiate and produce hemagglutinin-specific antibodies on-chip in response to a single dose of quadrivalent influenza vaccine

To evaluate if the LToC model supports B cell maturation and antibody responses over extended culture periods, we performed a longitudinal study in which LToCs were exposed to a quadrivalent influenza vaccine (inactivated, split virus; VaxiGrip Tetra® 2022/2023) overnight directly after on-chip seeding and then kept under perfusion for up to four weeks (Fig. 4A). The B cell differentiation markers CD27 and CD38 were used to track B cell activation and differentiation. While the naïve B cell fraction (CD27^-^CD38^-^) in unstimulated LToCs remained remarkably stable over time, influenza vaccine induced a gradual decrease in naïve B cell proportions, starting approximately seven days after treatment (Fig. 4B). Accordingly, the fraction of pre-germinal center (CD27^-^CD38^+^) and germinal center (CD27^+^CD38^+^) phenotype B cells gradually increased between weeks one and three of culture. Influenza vaccine further induced B cell differentiation into plasmablasts (CD27^+^CD38^+++^), starting as early as day 7. In line with the phenotypic B cell changes, vaccinated LToC produced higher concentrations of influenza hemagglutinin (HA)-specific antibodies than untreated LToC controls (Fig. 4C). Of note, the kinetics of HA-specific antibody production of two different donors tested revealed distinct immune regression times, with one donor only weakly and transiently responding, and the other producing sustained, high-magnitude antibody responses from week one to four. Influenza multiplex immunoassay analysis of the effluent from one experiment revealed that IgG antibodies in the effluent from vaccinated LToC bound to HA antigens from the four vaccine influenza H1N1, H3N2 and B strains, but also displayed reactivity against HA antigens from 13 related and unrelated strains and, to a smaller degree, four neuraminidase (NA) and five nucleoprotein (NP) antigens (Fig. 4D). In summary, the LToC enabled unprecedented retention of naïve B cells over four weeks in culture and the observed shifts in B cell differentiation and donor-specific HA-specific antibody responses suggest that the LToC model can successfully emulate individual immune responses against vaccines.

**Fig. 4:**
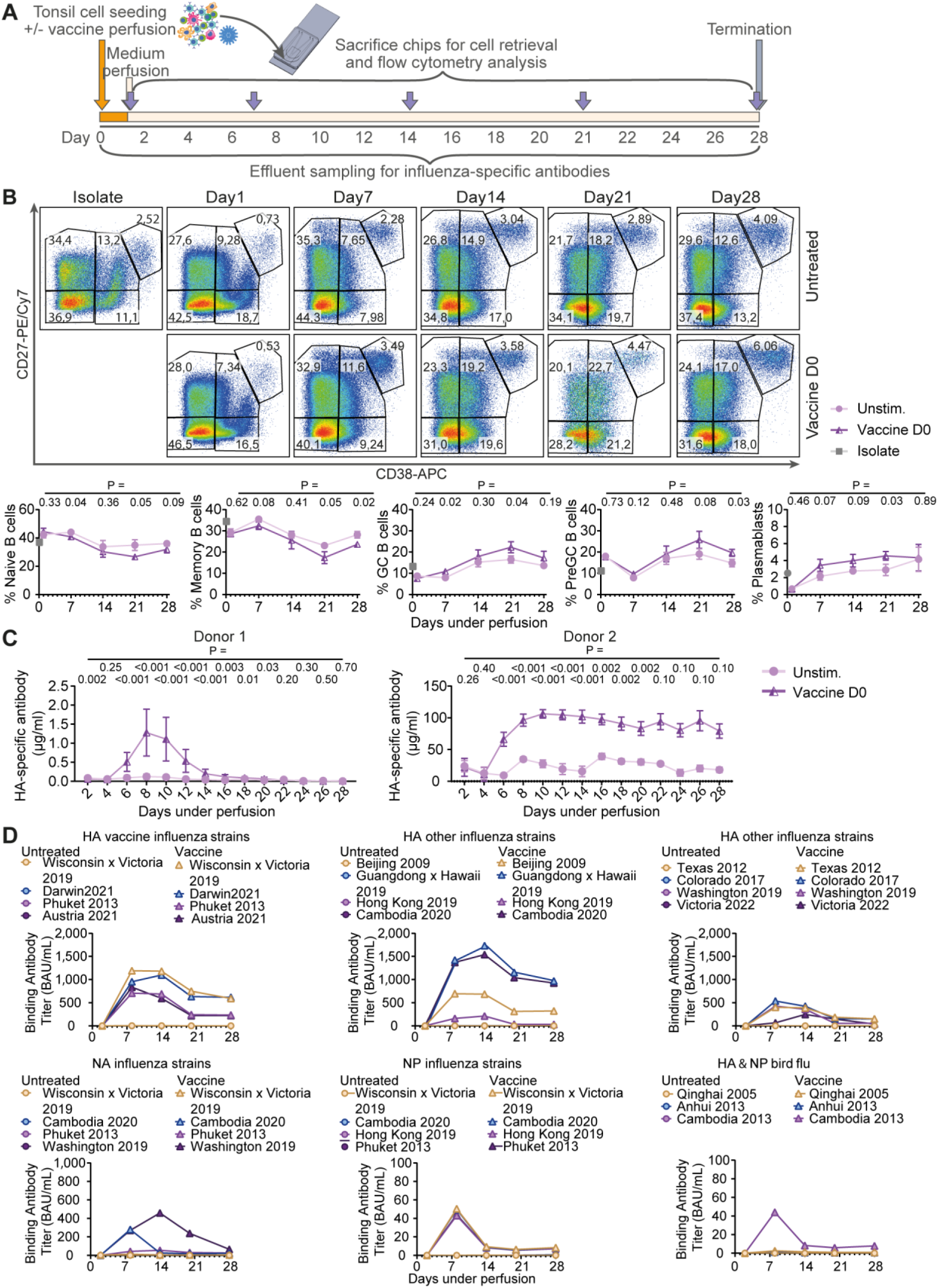
B cell differentiation and antibody production in LToC in response to inactivated split influenza vaccine. **A** Experimental timeline: LToCs were generated and perfused with medium (unstim.) or with vaccine-supplemented medium (vaccine D0) for one day, followed by perfusion with medium. Effluents were collected every other day and LToCs from each group were terminated for cell retrieval and flow cytometric analysis at different timepoints. **B** Flow cytometric analysis of B cell differentiation status within the B cell (CD19^+^) fraction. Exemplary flow cytometry plots (top) and graphs (bottom) showing percentages of naïve B cells (CD27^-^/CD38^-^), pre-germinal center B cells (CD27^-^/CD38^+^), germinal center B cells (CD27^+^/CD38^+^) plasmablasts (CD27^+^/CD38^+++^) and memory B cells (CD27^+^/CD38^-^) in thawed tonsil cell isolates and throughout LToC culture. n = 3 chips per condition. P-values were determined using multiple unpaired t test with Welch correction, without multiple comparison correction. **C** Quantification of influenza hemagglutinin (HA)-specific antibody release into effluent at different timepoints. n = 3-15 chips per condition (two donors). P-values were determined using multiple Mann-Whitney test, without multiple comparison correction. **D** Multiplex hemagglutinin subtype-specific antibody detection was applied to detect binding antibody units (BAU) per ml of antibodies against different influenza strains in effluents from donor 2. n = 3 chips per condition, standard deviation not shown.

### LToC remains responsive to vaccination for at least three weeks

To determine if untreated LToCs not only survive and persist in a predominantly naïve state, but also retain their ability to functionally respond to vaccine antigens, we next tested their ability to produce an antigen-specific response when vaccine was introduced shortly after culture initiation (day 0 or 2) or after three weeks of LToC culture. To monitor responsiveness, we perfused LToCs overnight with influenza vaccine on days zero, two, or 21, and monitored HA-specific antibody secretion for the following week. We observed a gradual increase in HA-specific antibody production in LToC vaccinated two days and three weeks after initiation of perfusion (Fig. 5). The HA-specific antibody concentrations in LToC effluents that were vaccinated two days and three weeks after culture were lower than the HA-specific antibody concentrations in LToC that received the vaccine dose directly at initiation of LToC perfusion, which might be caused by loss of some HA-specific B and T cells during culture. Nevertheless, these findings suggest that the untreated LToC remain functionally capable of responding to influenza vaccine with antibody secretion over at least three weeks of culture.

**Fig. 5:**
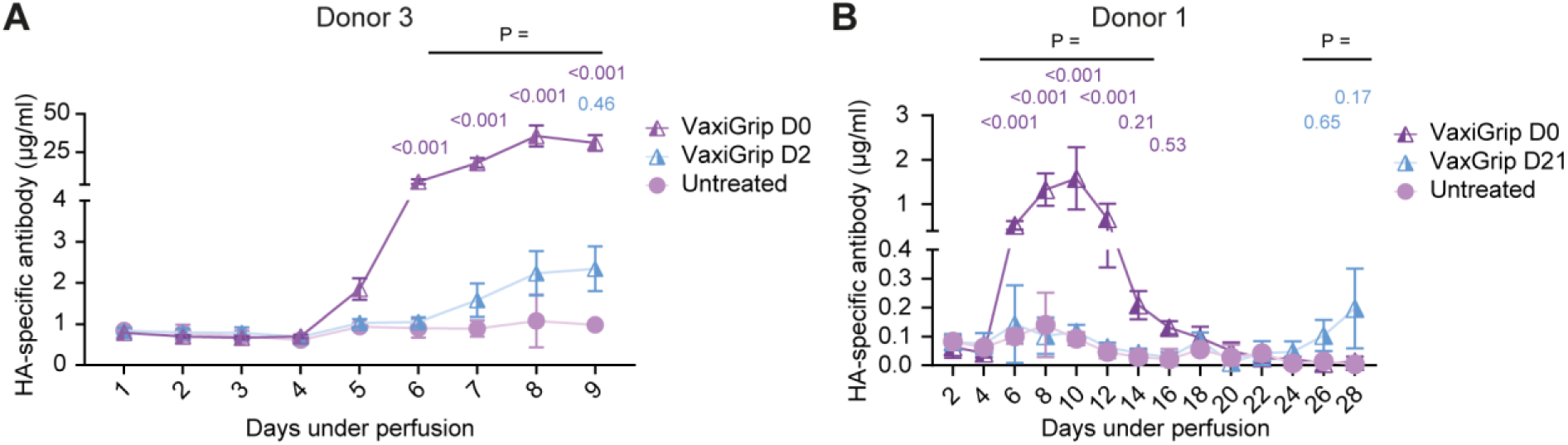
Longitudinal response to inactivated split influenza vaccine. Quantification of influenza hemagglutinin-specific antibody release into perfusion at different timepoints. LToCs were vaccinated at different timepoint of culture and medium was collected daily (**A**) or every other day (**B**). **A** unstim.: medium only, vaccine D0: vaccine from day 0 to day 1, vaccine D2: vaccine from day 2 to day 3. n = 3-4 chips per condition. **B** unstim.: medium only, vaccine D0: vaccine from day 0 to day 1, vaccine D21: vaccine from day 21 to day 22. n = 3 chips per condition. P-values were calculated using Ordinary two-way ANOVA with Dunnett’s multiple comparison test.

### Peripherally-derived antigen-presenting cells can migrate into LToC and modulate the extent and quality of humoral responses

Recent evidence suggests that the mode of vaccine administration influences the immune response in secondary lymphoid organs and consequently decides whether a vaccine induces durable and protective memory (*27*, *28*). To determine if the LToC model can be used to study immune response differences depending on the route of vaccine administration, we evaluated if the LToC can recruit antigen-presenting cells from perfused medium and if the direct addition of vaccine, the indirect addition of vaccine via antigen-presenting cells, or a combination of both can affect the downstream response in LToC. Therefore, we first isolated monocytes from autologous peripheral blood, differentiated them into monocyte-derived dendritic cells (moDCs), and generated LToCs two days before introducing quadrivalent influenza vaccine either through vaccine-primed moDCs (pre-vaccinated for three hours), by direct addition to the LToC culture medium as usual or by combining vaccine-primed moDCs and direct addition to the medium (Fig. 6A). To trace their migration, moDCs were labeled with CellTracker® DeepRed. Then we perfused LToC either with non-primed moDCs, pre-vaccinated moDCs, pre-vaccinated moDCs and soluble vaccine, the vaccine only, or left LToC untreated overnight. Seven days after the overnight treatment and continued perfusion with medium, labeled moDCs were detected within LToC, which demonstrated that the LToC induced active transmigration of moDCs from the medium through the membrane into the tissue chamber (Fig. 6B,C). Flow cytometry data from cells retrieved from LToC seven days after treatment showed that direct vaccination resulted in the previously described drop in naïve and memory B cells as well as the augmented proportions of pre-germinal center B cells, germinal center B cells and plasmablasts within the B cell compartment compared to untreated LToC, reminiscent of an immune response against influenza vaccine (Fig. 6D). Combined treatment with vaccine and pre-vaccinated moDCs induced similar but less strong trends in B cell differentiation as direct vaccination except for plasmablasts. The observed trends were even lower in LToC which were only perfused with pre-vaccinated moDCs. As expected, untreated moDCs had no impact on B cell differentiation status in LToC compared to untreated controls. In line with the flow cytometry data, measurements of HA-specific antibodies in the effluent from LToC revealed that direct vaccination resulted in the strongest antibody responses in two out of three donors, followed by weaker or equivalent antibody responses with combined direct and indirect vaccination, and the lowest but still detectable antibody responses by indirect vaccination alone (Fig. 6E). Results from influenza multiplex immunoassay analysis further showed that LToCs vaccinated with a combination of soluble vaccine and pre-vaccinated moDC produced less IgG antibodies targeting NP and NA influenza antigens and preferentially released IgGs against HA antigens from recent influenza strains (Fig. S2).

**Fig. 6:**
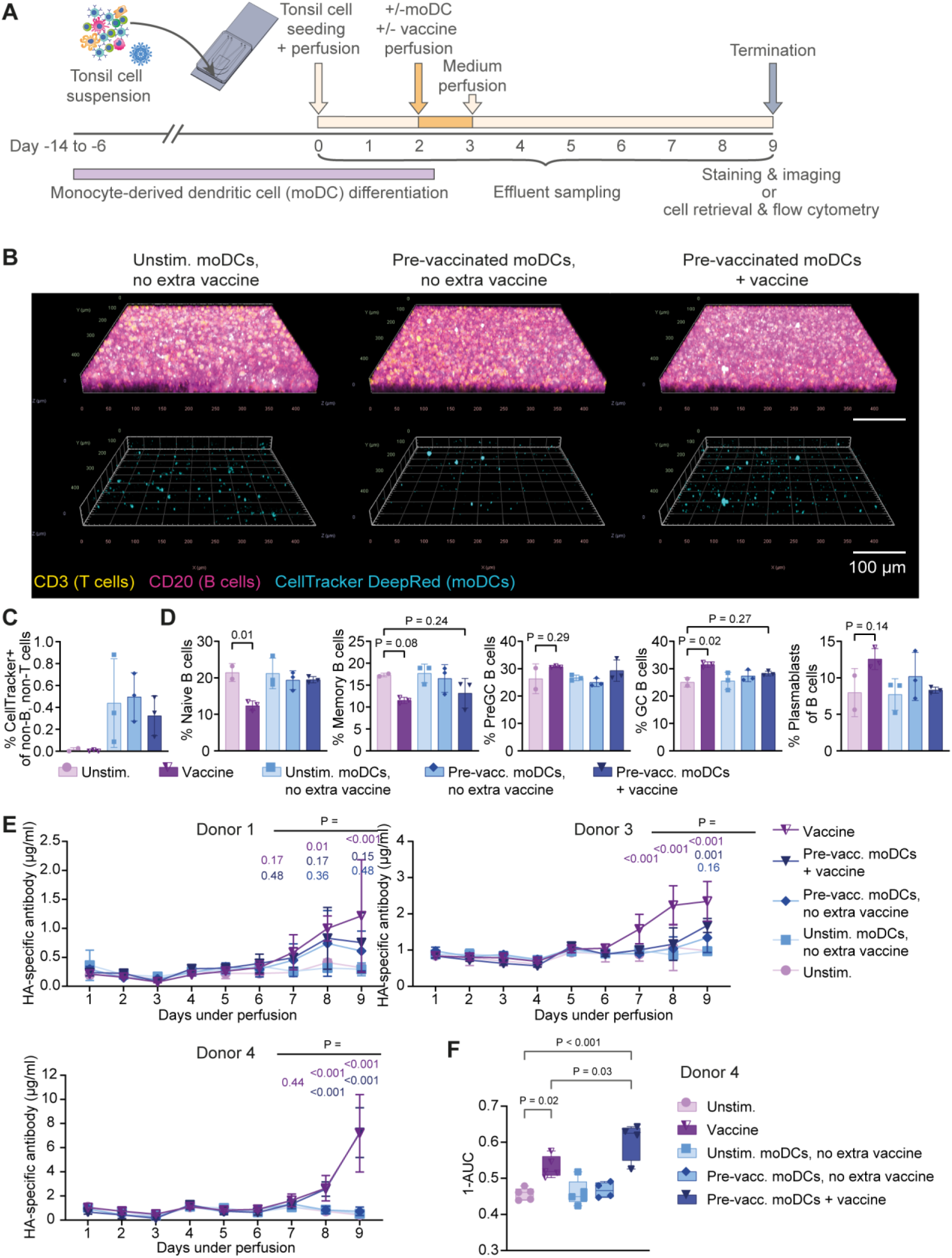
Impact of mode of antigen delivery on B cell responses in LToC. **A** Experimental timeline: Monocyte-derived dendritic cells (moDCs) were differentiated from thawed autologous peripheral blood-derived monocytes for at least six days and labelled with CellTracker before perfusion. LToCs were generated, perfused with medium for two days and then vaccinated overnight by supplementing perfused medium with either vaccine (vaccine D2), non-vaccinated moDC (unstim. moDCs, no extra vaccine), moDCs pre-vaccinated for three hours (pre-vacc. moDCs, no extra vaccine), both pre-vaccinated moDCs and vaccine (pre-vacc. moDCs + vaccine) or no vaccine (unstim.). Chips were stopped nine days after perfusion for analysis. **B** Exemplary 3D images of CellTracker-labelled moDCs, which transmigrated from the medium flow into the LToC. **C** Percentage of CellTracker-labelled non-B/non-T cells within cells retrieved from LToC analyzed by Flow cytometry. n = 2-3 chips per condition. **D** Percentages of naïve B cells (CD27^-^/CD38^-^), pre-germinal center B cells (CD27^-^/CD38^+^), germinal center B cells (CD27^+^/CD38^+^) plasmablasts (CD27^+^/CD38^+++^) and memory B cells (CD27^+^/CD38^-^) within B cells determined by Flow cytometric analysis. n = 2-3 chips per condition. **E** Quantification of influenza hemagglutinin-specific antibody release into effluent at different timepoints from three different donors. n = 2-5 chips per condition and donor. **F** Microneutralization assay using effluents from LToCs. Quantification by one minus area under the curve (1-AUC). All P-values were calculated using Ordinary two-way ANOVA with Dunnett’s multiple comparison test.

Finally, we performed a microneutralization assay with the effluent to assess whether the mode of antigen delivery affects the quality of the secreted antibodies. The neutralization capacity of antibodies from LToC that received combined vaccination clearly surpassed the neutralization of capacity of antibodies of LToC that were perfused with vaccine alone (Fig. 6F), despite similar magnitudes of HA-specific antibody (Fig. 6E). This suggests that the combination of full antigen and presentation of processed antigens via moDCs improved the quality of humoral responses in the LToC. In summary, these data show that LToC can actively recruit immune cells from the periphery and can be used to determine impact of the mode of antigen delivery on the strength and quality of immune response.

## DISCUSSION

Currently available human lymphoid tissue models are restricted by limited longevity, low cellular complexity, or constraints in the mode of antigen delivery. As a result, studies on the duration of antibody responses, effects of repeated vaccine dosing and further lymph node functions in health and disease have been mostly confined to animal studies. These limitations are addressed by the LToC reported in this work. The LToCs displayed exceptional long-term viability and functionality of human lymphoid tissue-derived cells, supported antigen-mediated B cell maturation and antibody responses and allowed to discern the magnitude and quality of antibody responses to direct and indirect vaccination modalities. The LToC model showcases how the fusion of appropriate cell sources and new organ-on-chip technology can advance modelling of physiological relevant processes towards applications such as vaccine testing and beyond.

The developed LT chip design ensured injection of highly consistent cell numbers, which is essential for reproducible immune responses between chips. The unique strategy of centrifugal loading introduced a compactness of the loaded tonsil cells that closely resembled the densities observed in native lymphoid tissue (*29–31*). The generated LToCs were marked by a stable cell density as well as stromal and immune cell composition throughout four weeks of culture. Appropriate cell density and correct composition of cellular subsets as provided by tonsil-derived cells is crucial for the preservation of central immune cell functions, as exemplified by the role of MHC self-recognition (*32*) and the dendritic cell compartment (*33*) in maintaining T cell antigen responsiveness in the absence of their cognate antigen in the lymph node. In particular the in vitro maintenance and differentiation of human B cell monocultures requires artificial signals such as addition of CD40 ligand and a mix of cytokines (*34*) or modified feeder cells (*35*), which, however, profoundly influence the non-specific activation and proliferation of B cells. Such activated cultured B cells do not entirely reflect germinal center processes in vivo and prevent comparisons of treatment effect on B cell subsets within a physiological context. Even in the previously published tonsil aggregates, the ratio of naïve B cells gradually dropped during three weeks of culture (*20*), indicating either an impaired survival of naïve B cells or a gradual collateral activation through accumulation of cytokines and waste products in static transwell or plate culture. The observed durability of B cells including naïve B cells in LToC was therefore a major advance and enables long-term interaction studies that require stable unstimulated controls beyond the scope of vaccination and re-vaccination studies.

Multiplexed analysis of lymphoid tissue-associated cytokines, general immune-related cytokines, complement components, extracellular matrix proteins, soluble Fas and soluble cytokine receptor isoforms revealed that LToC continuously released lymphoid-associated, chemotactic cytokines into the effluent including CXCL13 and CCL19, interleukins with chemotactic and immunomodulating functions and Collagen IVα1. In secondary lymphoid organs, CXCL13 is majorly produced by follicular dendritic cells (*36*, *37*), but can also be secreted by dendritic cells (*37*) and T follicular helper cells (*8*), while CCL19 is produced by stromal cells (*38*, *39*). Therefore, the secretion of signal molecules such as CXCL13 and CCL19 indicates that LToC not only preserved cell ratios but also supports a persisting homeostatic signaling of stromal and immune cells. Collagen type IV has been found along the reticular network of human (*40*) and non-human primate (*41*) lymph nodes, which is crucial for the trafficking of immune cells within the lymph nodes (*42*). The Collagen IVα1 found in the LToC effluents implies that the LToC is able to produce own extracellular matrix further enhancing functionality of this model.

The microfluidic LToC system did not impair the previously described self-aggregation of tonsil-derived cells(*20*), as the LToC displayed optically discernable areas of higher and lower density even within the compactly seeded tissue chambers, suggesting a similar spatial distribution of cells in LToC and tonsil aggregates. The LToC further allowed retrieval of cells and enabled in-depth analysis of phenotypic changes of cells during immune responses using flow cytometry. This analysis revealed that B cells from vaccinated LToCs undergo expected B cell differentiation processes consistent with the adaptive immune responses observed in the tonsil organoids (*20*, *43*), and therefore reflect features of an in vivo immune response. In line with the statically cultured tonsil aggregates, but in contrast to a previously published organ-on-chip model of ectopic lymphoid follicles built from blood-derived B and T cells and moDCs (*44*), vaccination with a quadrivalent influenza vaccine alone induced or re-activated HA-specific antibody responses in LToC generated from different patient material. This finding highlights the importance of using the full cell subset diversity found in lymphoid tissue, including stromal cells, to reliably model central immune responses. The magnitude and kinetics of the HA-specific antibody responses from LToC generated from different tonsil donors varied, which reflects the diversity of immune responses among individuals. Influenza multiplex immunoassay analysis to determine antibody binding of vaccine-targeted antigens vs. other influenza antigens showed that vaccination with inactivated split virus led to production of IgG antibodies which bound to all introduced H1N1, H3N2 and influenza B HA antigens, but also showed reactivity to the HA antigens from different strains that occurred between 2012 and 2022. LToCs further generated low antibody responses against influenza NA and NP antigens. This aligns with the finding that individuals vaccinated with different types of inactivated influenza HA vaccines largely raise cross-reactive or memory HA-specific antibody responses, but also low-scale antibody responses against NA and other influenza antigens (*45*).

Given the role of chemokines such as CXCL13 and CCL19 in immune cell attraction into the lymph node (*8*, *36–39*), the finding that the LToC produced multiple chemokines suggested that the LToC can attract immune cells from the perfused medium. Indeed, untreated LToC successfully recruited both untreated and pre-vaccinated moDCs from perfused medium into the tissue chamber. Although the LToC remains a single organ model, we demonstrated that the direct, indirect and combined vaccination strategies elicited immune responses of different magnitude and quality. The strongest effects on B cell differentiation and the highest HA-specific antibody production were observed when LToCs were directly exposed to the soluble vaccine antigens. Interestingly, the combined delivery of undigested vaccine and pre-vaccinated moDCs induced lower levels of B cell differentiation, raised lower HA-specific antibody responses in two out of three donors, produced less influenza NA- and NP-specific antibodies and resulted in production of antibodies with a superior neutralization capacity. This observation indicates that antigen-specific B cells in the LToC undergo a longer but more efficient and antigen-targeted antibody maturation if the antigen is delivered both in processed and unprocessed form. LToCs perfused with pre-vaccinated moDCs alone did not display observable changes on B phenotypical level, but still mounted a low HA-specific antibody response. Although HA-specific B cells require recognition of soluble vaccine to become activated (*4*, *7*), it has been shown that both resident and recently migrated dendritic cells can still display a fraction of unprocessed antigen on their surface and thereby promote T cell-dependent antibody responses in B cells in the lymph node (*46–48*). Therefore, pre-vaccinated moDCs might have activated low level B cell responses in the LToC via intact HA proteins on their surface, which were not entirely washed off prior to perfusion. Overall, differences in the magnitude of B cell differentiation and antibody responses dependent on the mode of antigen delivery demonstrated that the LToC can be used to study immune responses in a more physiological setting and to prevent overestimation of vaccine effect using human in vitro models. In a next step, the LToC model would benefit from integration of blood and lymphatic endothelial cell linings in the channel to better model vaccine delivery via lymph and blood and to model the next layer of interaction at the lymphatics and high endothelial venule site of lymphoid organs. This, however, poses specific challenges given the requirement of autologous endothelial cell sources to avoid the development of allogenic reactions against foreign cells in long-term LToC co-cultures.

To conclude, the LToC model presents a new tool with enhanced longevity to evaluate differences in immune responses to vaccines dependent on the mode of antigen delivery by comparing the germinal center induction in the lymphoid tissue, the quality of antibody responses and the level of cytokine production in perfused medium. Thereby, it can help to determine the efficacy of vaccine candidates in a more physiological setting across groups of individuals of different sex, age, and ethnicity. Beyond the scope of vaccine testing, the LToC model can further be used to study interactions between lymphoid tissue-derived human cells during homeostasis, immune responses and immunomodulatory therapies over the course of several weeks.

## MATERIALS AND METHODS

### Lymphoid tissue chip design and fabrication

Plastic film photomasks (Filmbelichtung24), depicting four tissue and medium layers of the lymphoid tissue (LT) chip, were drafted using AutoCAD (Autodesk) and used to fabricate SU-8 photoresist (Microresist Technologies) features on silicon wafers (Siegert Wafer) via photolithographic processes as described before (*22*, *49*). As visualized with SolidWorks (Dassault Sytèmes) (Fig. 2A), the SU-8 structures on the tissue layer wafer had three heights: 315 μm for the tissue chamber, 215 μm for the tissue seeding channel and 15 μm for the membrane inlay (area 9x18 mm). Medium channel structures on the medium layer wafer were produced at 200 μm height. Tissue and medium layers were fabricated via replica molding of polydimethylsiloxane (PDMS; Sylgard 184, Dow Corning). PDMS was homogeneously mixed in a 10:1 elastomer base to curing agent mass ratio and degassed in a desiccator. Medium layers were produced using a standard molding approach by pouring the PDMS mixture onto the silicon wafer master mold to obtain PDMS pieces with medium channel structures, followed by curing at 60°C for 4 h. PDMS pieces were cut to the size of the chip and ports were opened using a disposable biopsy punch (diameter 0.75 mm, World Precision Instruments). Tissue layers were exclusion molded by pouring PDMS solution onto the silicon wafer master mold and clamping it against a 5 mm-thick PMMA disk to produce a layer with through-hole tissue chamber structures (*49*). PDMS was cured at 60°C for 2 h. Before bonding, media layers and isoporous, polyethylene terephthalate (PET) membranes (9 mm x 18 mm, 5 μm pores, 60’000 pores/cm^2^; TRAKETCH PET 5.0 p S210 3 300, SABEU GmbH & Co. KG) featuring an ultra-thin glass-like coating (as described previously (*49*)) were cleaned with isopropanol and blow-dried with an air pistol and precision wipes (Kimtech Science) if necessary. The microfluidic chips were assembled in three bonding steps: 1) tissue layer to glass coverslip (for imaging, 24x40 mm, Knittel) or glass microscope slide (for cell retrieval, 76×26 mm, Menzel), 2) coated PET membrane to tissue layer-glass sandwich, and 3) medium layer to the sandwich from step 2). Bonding was achieved by oxygen plasma activation (75 W, 0.2 sccm O_2_, Diener Zepto, Diener Electronic) for 24 s before each assembly step, followed by incubation at 60°C for at least 30 min. After step 3, assembled chips were baked at 60°C overnight.

### Consent and human tissue and blood sample collection

One half of each tonsil and blood from four adult donors with recurrent tonsillitis were collected at the Head and Neck Surgery of the University Clinic Tübingen after informed consent as approved by the Ethical Committee of the Eberhard Karls University Tübingen (No. 346/2022BO2).

### Tonsil cell isolation and cryopreservation

Tonsil biopsies were transferred into decontamination solution containing Ham’s F12 medium (HyClone, Cytiva) supplemented with 2% FetalClone II Serum (FCS) (HyClone, Cytiva), 2X antibiotic-antimycotic solution (Gibco) and 1X Normocin (Invivogen) directly after resection and incubated for at least 30 min at 4°C. Tonsil tissue was washed in Ham’s F12 medium with 2% FCS and mechanically dissociated as described before (*20*). Gradient centrifugation (Histopaque 1077, Sigma-Aldrich) was applied to remove debris and tonsil-derived cells were cryopreserved in FCS with 10% dimethyl sulfoxide (DMSO) (PanReac AppliChem) using a CoolCell^TM^ Cell Freezing Container (Corning).

### Isolation and cryopreservation of blood monocyte

Blood was mixed 1:1 with phosphate buffered saline without calcium and magnesium (PBS-) (PAN-Biotech) and separated using Histopaque 1077 and SepMate^TM^-50 (IVD) (STEMCELL Technologies) for density centrifugation (1,200 xg, 12 min) within one to five hours after blood collection. After centrifugation, peripheral blood mononuclear cells (PBMCs) were poured into a fresh tube and washed twice in PBS− supplemented with 0.1% BSA (Sigma) and 2 mM EDTA (Gibco). PBMCs were used directly for monocyte isolation using Pan Monocyte Isolation Kit, human, LS columns and QuadroMACS (all Miltenyi Biotec) according to manufacturer’s protocol. Monocytes were frozen in RPMI 1640 medium (Gibco) with 20 % FCS and 10 % DMSO using a CoolCell^TM^ Cell Freezing Container (Corning).

### Tonsil cell seeding on-chip

Chips were sterilized and hydrophilized with O_2_-plasma (75 W, 0.2 sccm O_2_, Diener Zepto, Diener Electronic) for 5 min. Then channels were entirely flushed through one tissue channel port with 70% ethanol using 200 μL ultrapoint pipette tips (Thermo Fisher Scientific), washed with PBS- and stored with PBS- (PAN-Biotech) at room temperature (RT) until use. Cryopreserved cells were thawed in thaw medium (RPMI 1640 with GlutaMAX supplement (Gibco) with 10% FCS and 1X antibiotic-antimycotic), resuspended to 3.75 x 10^6^ cells/mL in complete medium (RPMI 1640 with GlutaMAX, 10% FCS, 1X nonessential amino acids (Gibco), 1X sodium pyruvate (Gibco), 1X antibiotic-antimycotic, 1X Normocin and 1X Insulin-Transferrin-Selen (ITS-G) (Gibco)) and stored on ice.

For chip loading one tissue channel port was closed with a sterilized metal wire with 0.7 mm diameter (Menzanium). 8 μL of cell suspension (= 3 x 10^6^ cells) was injected into the other tissue channel port and the second tissue channel port and both medium channel ports were closed. Loaded chips were transferred into 50 mL tube (Greiner Bio-One) with the tissue chamber facing the bottom of the tube and centrifuged at 100 xg for 3 min. After centrifugation, the tissue channel ports were opened, the tissue channel was flushed with 12 μl 3-D Life Dextran-CD Hydrogel SG (Cellendes) using 4 mM CD-linker and the tissue channel ports were closed again. Finally, the medium channel was opened and flushed with 100 μL complete medium by hydrostatic pressure by attaching one empty 200 μL ultrapoint pipette tip at one port and inserting one medium-filled tip to the other port. The gel was allowed to crosslink at 37°C, 5% CO_2_ and 95% rH for at least 30 min.

### Perfused chip culture, on-chip vaccination and effluent collection

For chip perfusion, Tygon tubings (0.51 mm inner diameter, Tygon ND 100-80 Medical Tubing, Saint-Gobain Performance Plastics Pampus GmbH) were connected to a 21 GA stainless steel plastic hub dispensing needle (KDS2112P, Weller Tools GmbH) at one side and to a blunt 21 GA stainless steel needle (detached from the plastic hub by dissolving the glue overnight in a 70% ethanol solution) on the other side and autoclaved. The plastic hub dispensing needle was attached to a syringe (B. Braun) filled with complete medium with 0.1 μg/mL recombinant human BAFF (Biolegend), inserted into a syringe pump (LA-190, Landgraf Laborsysteme HLL GmbH), and the blunt needle was inserted into one medium channel port. The other port was connected to a blunt 21 GA needle attached to a Tygon tubing inserted into an effluent collection tube. The chips were perfused with 40 μL/h in push mode at 37°C, 5% CO_2_ and 95% rH. For on-chip vaccination, complete medium with 0.1 μg/mL recombinant human BAFF supplemented with 1:2,000 diluted (= 7.5 ng/ml HA per strain) quadrivalent influenza vaccine (inactivated, split virus; VaxiGrip Tetra® 2022/2023, 15 μg/mL HA per strain, Sanofi Pasteur) was perfused overnight (∼16 h) and then changed to complete medium with 0.1 μg/mL recombinant human BAFF. Medium was changed every three to four days.

Depending on the experiment, effluent samples from each chip were collected every day or every other day, centrifuged at 10,000 xg for 3 min to remove debris. For multiplex cytokine assay and antibody measurements samples were stored at -80°C until further analysis. For lactate dehydrogenase assay, effluents were diluted 1:10 in LDH storage buffer (200mM Tris-HCl (pH 7.3) (Sigma-Aldrich), 10% Glycerol (Carl Roth), 1% BSA) and stored at -20 to -80°C.

### In vitro differentiation of monocyte-derived dendritic cells (moDCs)

Monocyte-derived dendritic cell differentiation protocol was adapted from previously published protocols (*50*, *51*). Cryopreserved monocytes were thawed in X-VIVO^TM^ 15 medium (Lonza), plated at 2-3 x 10^6^ cells/well into a 6-well plate (Greiner Bio-One) in dendritic cell (DC) differentiation medium (X-Vivo^TM^ medium supplemented with 10% human serum (Sigma-Aldrich), 1% penicillin/streptomycin (10,000 U/ml, Pan Biotech), 500 IU/mL GM-CSF and 500 IU/mL IL-4 (both premium grade, Miltenyi Biotec)) and incubated at 37°C, 5% CO_2_ and 95% rH. The next day, the medium was removed and replaced with fresh DC differentiation medium. 50% medium change was performed every two to three days until harvesting. Cells were differentiated for a minimum of six days.

### CellTracker-labelling, pre-vaccination and perfusion of monocyte-derived dendritic cells

Autologous moDCs were harvested, washed in PBS with calcium and magnesium (PBS+), resuspended in 1:1,500 (for imaging) or 1:20,000 (for flow cytometry analysis) diluted CellTracker Deep Red (Invitrogen) in PBS+ (Pan Biotech) (= 0.67 μM final concentration) and incubated at 37°C, 5% CO_2_ and 95% rH for 15 min. Cells were then washed and resuspended to 2 x 10^6^ cells/mL in complete medium. Cells were split, mixed 1:1 either with complete medium without (unstimulated moDCs) or with (pre-vaccinated moDCs) 1,1000 diluted quadrivalent influenza vaccine and incubated at 37°C, 5% CO_2_ and 95% rH. After three hours of incubation, unstimulated and pre-vaccinated moDCs were washed once with complete medium and resuspended to 1 x 10^6^ cells/mL in complete medium with BAFF.

The perfused chip setup was removed from the incubator, the tubing connecting the medium channel to the effluent collection tube was removed and replaced with a 200 μL ultrapoint tip containing 50 μL complete medium with BAFF. Negative pressure was introduced by running the attached syringe pump setting at 40 μL/h in pull mode for at least 30 min 37°C, 5% CO_2_ and 95% rH. Under continued perfusion, 50 ul of untreated moDCs or pre-vaccinated moDCs (= 50,000 cells/chip) or complete medium with BAFF was added to the pipette tips of respective chips. For unstimulated LToC, LToC with unstimulated moDCs without extra vaccine and LToC with pre-vaccinated moDCs without extra vaccine, 350 ul of complete medium with BAFF was added to the respective tips. For LToC conditions with vaccine only or pre-vaccinated moDCs with vaccine, 350 ul of complete medium with BAFF and 1:2’000 diluted vaccine was added. Chips were perfused at 20 μL/h in pull mode overnight (∼16 h) at 37°C, 5% CO_2_ and 95% rH and changed back to perfusion with complete medium with BAFF at 40 μL/h in push mode at 37°C, 5% CO_2_ and 95% rH. Medium was changed once more after three to four days of perfusion.

### Immunofluorescence staining on-chip

LToC was washed with 100 μl PBS+ through the medium channel by applying hydrostatic pressure using 200 μL ultrapoint pipette tips as described above. LToC was then fixed with ROTI Histofix 4% formaldehyde (Carl Roth) for 15 min or overnight at RT on a rocker (20 rpm, IKA). Then LToCs were flushed once with PBS-, washed twice with PBS- for 15 min on a rocker, blocked with 3% BSA in PBS- for 30 min at RT and incubated with 1:25-1:50 diluted CD3 (rabbit, SP162, Abcam), CD20-PE (human, REA780, Miltenyi Biotec) and optionally with CD38-APC (mouse, HIT2, Biolegend) antibody in 0.3% BSA in PBS-at 4°C overnight or for two days. LToC was flushed once and washed twice with PBS-/0.01% Tween 20 (Sigma-Aldrich) for 15 min on a rocker protected from light. Finally, LToC was incubated with 1:50 diluted rabbit IgG (H+L) cross-absorbed secondary AF488-labelled antibody (goat, polyclonal, Invitrogen) in 0.3% BSA in PBS- for three hours to two days at 4°C in the dark, flushed once with PBS-, washed twice with PBS- for 15 min and once for two to three hours on a rocker and stored at 4°C with PBS- until image acquisition.

### Calcein AM and propidium iodide staining on chip

LToC was washed with 100 μl PBS+ through the medium channel by applying hydrostatic pressure using 200 μL ultrapoint pipette tips as described above and viable cells were stained with 2 µg/mL Calcein AM Green in PBS+ (Thermo Fisher Scientific) for 1 h at 37°C. Then, propidium iodide was diluted to 50 µg/mL in PBS+ and added for 5 min at 37 °C for labeling nuclei of dead cells. Samples were flushed once with PBS+ and washed with PBS+ three times for 15 min on a rocker. Stained chips were fixed with ROTI Histofix 4% formaldehyde overnight at RT on a rocker, washed three times with PBS- for 15 min on a rocker and stored at 4°C until image acquisition.

### Image acquisition

Images were obtained using a confocal Laser-Scanning-Microscope (LSM 880, Zeiss, Germany). Images were processed by adjusting brightness and contrast using ZEN lite Software (Version 3.4, Zeiss)

### Cell retrieval, flow cytometry staining and analysis

For cell retrieval, medium channel ports were closed with metal wires with 0.7 mm diameter and the PDMS layer above the tissue chamber of LToCs was punched out using a 6 mm biopsy puncher (pfm medical). The membrane was carefully lifted with a pipette tip with 100 μL complete medium, cells were resuspended and transferred into cell collection tubes on ice. The medium channel was flushed with 100 μL complete medium once from each port, medium was collected from the tissue chamber and transferred to respective collection tubes. Chips were incubated 20-30 min on ice with 100 μL FACS buffer (3% FCS, 2 mM EDTA, PBS-) then, resuspension and medium channel wash steps were repeated with FACS buffer and all suspensions were added to respective collection tubes. Cell collection tubes were then centrifuged, and cells were transferred into 96-well plate V-bottom plates for flow cytometry staining in reduced volume. Cells were washed once with FACS buffer, blocked with 1:25 diluted human Fc Block (BD Biosciences) for 10 min on ice and stained for 30 min on ice with live/dead Zombie Aqua Dye (1:200, Biolegend) and 2X antibody cocktail in FACS buffer to reach final concentrations of: CXCR5-FITC (1:33, J252D4), CD8-PerCP (1:50, HIT8a), CD38-APC (1:200, HIT2) for long-term experiment or CD38-BV785 (1:100, HIT2) for moDC perfusion experiment, CD45-AF700 (1:100, HI30), HLA-DR-BV711 (1:100, L243), CD3-BV605 (1:100, OKT3), CD4-BV650 (1:100, RPA-T4), CD19-PE (1:200, HIB19), CD27-PE/Cy7 (1:100, O323), IgD-APC/Cy7 (1:50, IA6-2) (all from Biolegend). Samples were then washed twice with FACS buffer, fixed with 4 % PFA (Carl Roth) in PBS- for 10 min at RT, washed twice with FACS buffer and stored at 4°C with FACS buffer until measurement with BD LSRFortessa flow cytometer (BD Biosciences). Dara was analyzed using FlowJo V10.10 software.

### Lactate dehydrogenase assay

Lactate dehydrogenase (LDH) release into the media was detected according to the manufacturer’s instructions using the LDH-Glo Cytotoxicity Assay kit (Promega). Luminescence was measured in duplicate in an opaque 384-well plate (Lumitrac, Greiner Bio-One) using the Tristar 5 plate reader (Berthold). LDH standard curves to determine LDH activity in each sample in mU/ml were calculated using GraphPad Prism 10.4.0 software.

### Multiplex cytokine assay

Cytokines were measured using a commercial custom multiplex human magnetic Luminex assay (R&D Systems, Wiesbaden, Germany, cat no. LXSAHM, see Table S1Table S1 Table S1). Effluent samples were analyzed according to manufacturer’s protocol. In brief, samples were diluted 1:2 in assay buffer (CD RD6-52) and pipetted into individual wells of a 96-well plate. Standards provided by the manufacturer were included on each plate in duplicate. Bead mixes were diluted in diluent buffer (RD2-1) according to manufacturer’s protocol and added to the diluted effluent. Plates were then sealed and incubated on a thermomixer (800 rpm, 21°C) for 2 hours, after which unbound sample was removed and beads were washed three times using an automated plate washer (Biotek Multiflo FX). 50 µL biotinylated detection antibody was added to each well and incubated at 800 rpm, 21°C for 1 hour. To remove unbound detection antibody, the plates were washed three times, after which 50 µL Streptavidin-PE conjugate was added to each well and incubated for a further 30 mins at 800 rpm, 21°C. The beads were then washed three times, resuspended in 100 µl of wash buffer and analysed using a FLEXMAP 3D running Luminex xPONENT (v. 4.2) software (Luminex, Austin, USA). Median fluorescence intensity (MFI) values were back-calculated using a 5PL regression fit to the standard samples (Bio-Plex Manager, version 6) to determine cytokine concentrations.

### Hemagglutinin-specific antibody detection by enzyme-linked immunosorbent assay (ELISA)

ELISA for detection of influenza hemagglutinin-specific antibodies was performed as previously described (*43*). In brief, EIA/RIA high bind microplates (Corning) were coated with 2 μg/ml quadrivalent influenza vaccine (inactivated, split virus; VaxiGrip Tetra® 2022/2023, Sanofi Pasteur) in 100 mM sodium carbonate/bicarbonate buffer (Sigma-Aldrich) overnight at 4°C and blocked with 1% BSA in PBS- for 2 h at RT. Effluents were diluted 1:10 or 1:100 in PBS- and added for 1 h at RT. For detection of bound antibodies, horseradish peroxidase-conjugated anti-human secondary antibodies to IgM/IgG/IgA (polyclonal, Abcam) were applied for 1 h at RT. Plates were developed to SureBlue TMB substrate (KPL), quenched with sulfuric acid (2N) (PanReac Applichem) and measured at 450 nm using the Tristar 5 plate reader (Berthold). Concentrations of HA-specific antibodies were estimated using a human monoclonal influenza A hemeagglutinin IgG1 anibody (H1N14-M, alpha diagnostic international) as a standard.

### Multiplex hemagglutinin subtype-specific antibody detection

Various influenza antigens (Table S2) were immobilized on spectrally distinct populations of MAGplex beads using EDC/sNHS coupling as previously described (*52*). Following coupling, beads were stored at 4°C, before being combined into a 25x mix prior to use. Effluent samples were thawed, diluted in assay buffer (*53*) with 25 µL then transferred into individual wells of a 96-half well plate, after which 25 µL of 1x Bead Mix was added. Samples were then incubated for 2 hours at 21°C, 750 rpm, after which unbound antibodies were removed by washing 3x with wash buffer (PBS + 0.05% Tween20). To detect bound IgG antibodies, RPE-conjugated human IgG (3 µg/mL) was added to each well and incubated at 21°C, 750 rpm for 45 mins. Unbound detection antibodies were removed by washing 3x with wash buffer. The beads were then resuspended in 100 µL Wash Buffer, shaken briefly for 3 mins at 750 rpm and then measured using an INTELLIFLEX-DRSE with the following settings: 80 µL sample volume, 50 beads per ID, and a gating range of 7000-17000. For conversion to BAU/mL, the 2nd International Standard (10/202) was included on each plate, with a 7-parameter non-linear regression used to calculate IU/mL. For quality control, intra-well IgG and gt-a-hu-IgG beads as well as plate control samples were included.

### Microneutralization assay

Microneutralization (MN) assays in this study were conducted with adaptations from previously outlined protocols (*54*). Madin-Darby canine kidney cells (MDCKs, ATCC) were cultivated in Eagle’s minimum essential medium (EMEM, ATCC) supplemented with 1% antibiotic-antimycotic and 10% heat-inactivated FBS and incubated at 37°C with 5% CO_2_. Only early passaged cells were used for MN assays and subculturing of cells occurred upon reaching 80–85% confluency. One day prior to the assay, MDCK cells were subcultured into flat-bottomed 96-well plates at a density of 1.1 × 10^5^ cells/ml in 100 µL (1.1 x 10^4^ cells/well). Organoid culture supernatants were prepared at 100 μl and diluted (1:5) in virus growth media (VGM) consisting of serum-free EMEM supplemented with 0.6% BSA (Sigma-Aldrich) and 1 µg/mL N-p-Tosyl-l-phenylalanine chloromethyl ketone (TPCK)-treated trypsin (Worthington Biochemical) and then serially diluted (two-fold) in VGM, 50 μl were left over in all wells. The infectious A/California/07/09 X A/Puerto Rico/8/1934 reassortant H1N1 virus (BEI NR-44004) was diluted to 50 TCID50 per 50 µL in VGM and subsequently added to the serially diluted supernatants, followed by an incubation period of 1 hour at 37°C and 5% CO_2_. Replicate control samples consisting of 100 μl diluted virus only or 100 μl VGM only were also prepared. After incubation, the media from MDCK monolayers was replaced with the serum-virus mixtures and further incubated for 1 hour. Following this, the serum-virus mixtures were substituted with 200 µL of VGM supplemented with 2% FBS, and cells were incubated for 48 hours at 37°C. Post-incubation, media was removed and cells were fixed in 4% paraformaldehyde (PFA) in PBS for 30 minutes, washed once in PBS, and then permeabilized in 0.1% PBS/Triton X-100 (PBS-T) at room temperature for 15 minutes. Cells were washed twice with PBS and blocked using 200 μl of blocking buffer (3% BSA) in PBS for 1 hour at room temperature. Influenza virus nucleoprotein (NP) was detected utilizing an equal mixture of anti-NP mAbs (Millipore Cat Nos. MAB 8257 and MAB 8258) diluted (1/1000) in blocking buffer, followed by horseradish peroxidase (HRP)-conjugated anti-mouse IgG (KPL) diluted at 1:3000 in blocking buffer. Plates were developed in TMB peroxidase substrate and reactions were halted using 1N HCl. Finally, assays were quantified in an ELISA plate reader at 450 nm using SoftMax Pro 7.1 software. The area under the curve (AUC) of titration curve was determined per condition, followed by 1 minus AUC (1-AUC) calculation.

### Statistical analysis

All statistical analyses were performed using GraphPad Prism 10.4.0 software. Before statistical analysis, Shapiro-Wilk test was performed to test for normality of distribution of results within same groups. Multiple unpaired t test with Welch correction without multiple comparison correction was applied for normally distributed results, multiple Mann-Whitney test without multiple comparison correction was used for non-normally distributed results and Ordinary two-way ANOVA with Dunnett’s multiple comparison test was performed for comparison of multiple groups against unstimulated control as indicated in figure legends. Unless otherwise specified, standard deviation was depicted in every graph.

## Acknowledgments

We thank all the patients who participated in this study. We further acknowledge the Flow Cytometry Core Facility Klinik Berg at the University of Tübingen for providing shared instruments for flow cytometric sample acquisition.

## Author contributions

E.J.S.B established the chip fabrication process, F.B, J.B. and S.B. provided tonsil tissue and blood, C.T., A.S., A.W.E., D.V. and L.C. fabricated chips and isolated cells, C.T., A.S., A.W.E. and D.V. performed cell culture, chip experiments, flow cytometry, imaging, LDH assays, HA-specific antibody ELISAs and related analysis and data presentation, Z.W.W. performed and analyzed microneutralization assays, A.D. and P. M. established muliplexed hemagglutinin subtype-specific antibody assay, A.D. designed multiplex cytokine assay, M.F. performed all multiplex assays and performed analysis together with A.D.. C.T., P.L. and L.E.W. designed the study. J.M. provided equipment and input for imaging and image data analysis. C.T. wrote the manuscript with support from P.L., L.W., A.D. and J.M.. This work was supervised by P.L. with input from L.E.W.. All authors have read the manuscript, provided input and agree with its submission.

