## Supplemental_Figures_and_Tables for "Lymphoid-tissue-on-chip recapitulates human antibody responses in vitro"

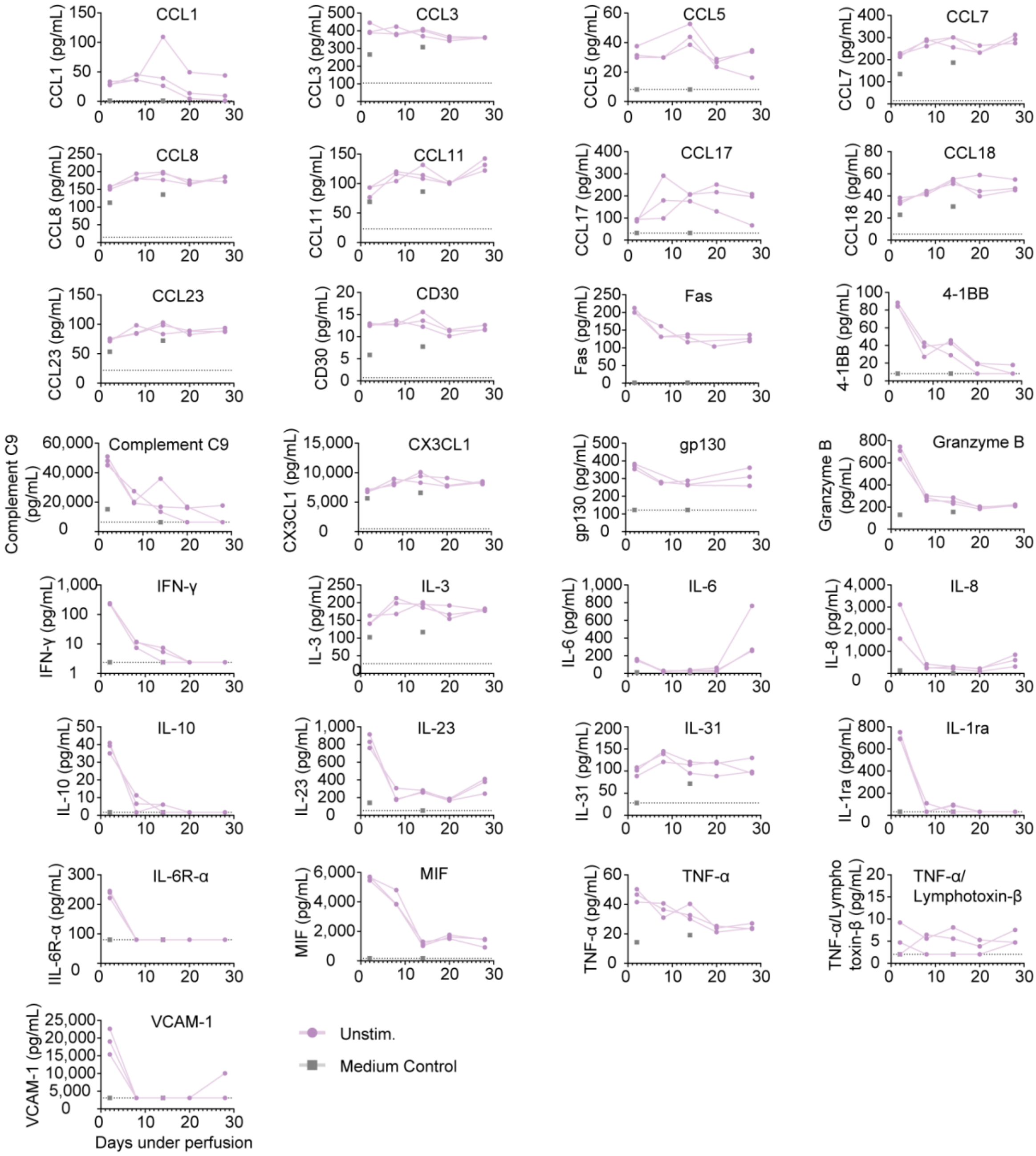


**Fig. S1 Cytokines and other signal molecules in LToC.** Multiplex cytokine analysis of LToC effluents (unstim.) and medium control. Effluents were collected every other day and effluents from day 2, 8, 14 and 28 were measured for secretion of respective analytes. Each continuous line depicts effluents from one LToC. Dashed lines indicate detection limit of respective analyte. Medium controls were measured from day 2 and day 14.

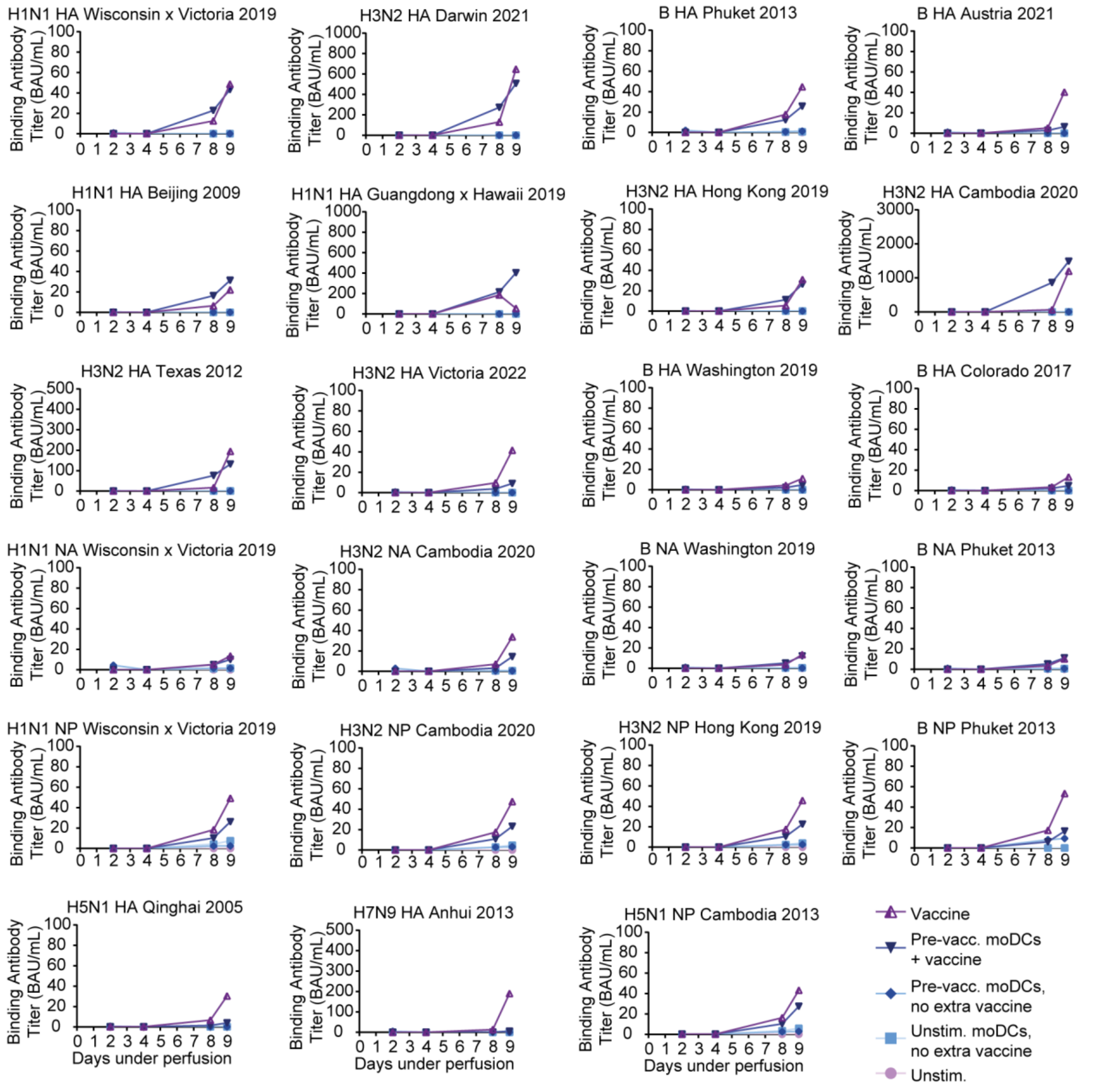


**Fig. S2 Impact of mode of antigen delivery on influenza antigen-specificity.** LToCs were generated, perfused with medium for two days and then vaccinated overnight by supplementing perfused medium with either vaccine (vaccine D2), non-vaccinated moDCs (unstim. moDCs, no extra vaccine), moDCs pre-vaccinated for three hours (pre-vacc. moDCs, no extra vaccine), both pre-vaccinated moDCs and vaccine (pre-vacc. moDCs + vaccine) or no vaccine (unstim.). Multiplex hemagglutinin subtype-specific antibody detection was applied to detect binding antibody units (BAU) per ml of antibodies against different influenza strains in effluents from donor 4. n = 2 chips per condition, standard deviation not shown.

| Bead Mix | Analyte |
|---|---|
| 1 | IL-1 beta/IL-1F2 |
| 1 | IL-4 |
| 1 | IL-2 |
| 1 | G-CSF |
| 1 | MBL |
| 1 | CCL17/TARC |
| 1 | IL-12/IL-23 p40 |
| 2 | BDNF |
| 2 | MIF |
| 2 | IL-27 |
| 2 | Pentraxin 3/TSG-14 |
| 2 | IL-17E/IL-25 |
| 2 | Prolactin |
| 2 | Lymphotoxin-alpha/TNF-beta |
| 2 | IL-13 |
| 2 | M-CSF |
| 2 | Myeloperoxidase/MPO |
| 2 | FGF-23 |
| 2 | RBP4/Retinol Binding Protein 4 |
| 2 | CXCL11/I-TAC |
| 2 | FGF acidic/FGF1 |
| 2 | BCMA/TNFRSF17 |
| 2 | TIMP-1 |
| 3 | TNF-alpha |
| 3 | gp130 |
| 3 | IL-3 |
| 3 | CCL7/MCP-3/MARC |
| 3 | IL-6R alpha |
| 3 | CXCL13/BLC/BCA-1 |
| 3 | CCL18/PARC |
| 3 | CCL3/MIP-1 alpha |
| 3 | IL-31 |
| 3 | CCL8/MCP-2 |
| 3 | CCL19/MIP-3 beta |
| 3 | BAFF/BLyS/TNFSF13B |
| 3 | CX3CL1/Fractalkine |
| 3 | Complement Component C9 |
| 3 | Resistin |
| 3 | IL-15 |
| 3 | Collagen IV alpha 1 |
| 3 | CCL23/MPIF-1 |
| 3 | Granzyme B |
| 3 | CD30/TNFRSF8 |
| 3 | IFN-alpha |
| 3 | Chemerin |
| 3 | IL-6 |
| 3 | Fas/TNFRSF6/CD95 |
| 3 | CCL5/RANTES |

| | |
|---|---|
| 3 | CCL11/Eotaxin |
| 4 | IL-28A/IFN-lambda 2 |
| 4 | 4-1BB/TNFRSF9/CD137 |
| 4 | IL-33 |
| 4 | CCL1/I-309/TCA-3 |
| 4 | Complement Component C5a |
| 4 | DPPIV/CD26 |
| 4 | IL-7 |
| 4 | IFN-beta |
| 4 | IL-10 |
| 4 | CCL2/JE/MCP-1 |
| 4 | Angiopoietin-2 |
| 4 | CXCL2/GRO beta/MIP-2/CINC-3 |
| 4 | CCL14/HCC-1/HCC-3 |
| 4 | IFN-gamma |
| 4 | IL-1ra/IL-1F3 |
| 4 | CCL20/MIP-3 alpha |
| 4 | HGF |
| 4 | CCL15/MIP-1 delta |
| 4 | CCL4/MIP-1 beta |
| 4 | IL-1 alpha/IL-1F1 |
| 4 | IL-17/IL-17A |
| 4 | APRIL/TNFSF13 |
| 4 | Adiponectin/Acrp30 |
| 4 | GM-CSF |
| 4 | FGF basic/FGF2/Bfgf |
| 4 | IL-8/CXCL8 |
| 4 | IL-5 |
| 4 | IL-12 p70 |
| 4 | VCAM-1/CD106 |
| 4 | VEGF-A |
| 4 | Complement Factor D/Adipsin |
| 4 | IL-21 |
| 4 | alpha-Fetoprotein/AFP |
| 4 | BMP-2 |
| 4 | BMP-4 |
| 4 | BMP-9 |
| 4 | Serpin A12 |
| 4 | Aggrecan |
| 4 | IL-2 |
| 4 | CXCL1/GRO alpha/KC/CINC-1 |
| 4 | IL-18/IL-1F4 |

**Table S1 List of analytes included within cytokine assay as well as their specific mixes.** Analytes were split into 4 bead mixes to avoid cross-reactivity.

| Subtype | Protein | Strain | Manufacturer | Catalogue # |
|---|---|---|---|---|
| H1N1 | HA | Wisconsin x Victoria 2019 | Sino Biological | 40787-V08H |
| H1N1 | HA | Beijing 2009 | Sino Biological | 40035-V08H |
| H1N1 | HA | Victoria 2022 | Sino Biological | 40938-V08H |
| H1N1 | HA | Guangdong x Hawaii 2019 | Sino Biological | 40717-V08H |
| H3N2 | HA | Cambodia 2020 | Sino Biological | 40789-V08H |
| H3N2 | HA | Hong Kong 2019 | Sino Biological | 40721-V08H |
| H3N2 | HA | Darwin 2021 | Sino Biological | 40859-V08H |
| H3N2 | HA | Texas 2012 | Sino Biological | 40354-V08H1 |
| Yamagata | HA | Phuket 2013 | Sino Biological | 40498-V08H1 |
| Victoria | HA | Austria 2021 | Sino Biological | 40862-V08H |
| Victoria | HA | Colorado 2017 | Sino Biological | 40581-V08H |
| Victoria | HA | Washington 2019 | Sino Biological | 40722-V08H |
| H7N9 | HA | Anhui 2013 | Sino Biological | 40103-V08H |
| H5N1 | HA | Qinghai 2005 | Sino Biological | 40117-V08B |
| H1N1 | NA | Wisconsin x Victoria 2019 | Sino Biological | 40785-V08B |
| H3N2 | NA | Cambodia 2020 | Sino Biological | 40784-V08B |
| Yamagata | NA | Phuket 2013 | Sino Biological | 40502-V07B |
| Victoria | NA | Washington 2019 | Sino Biological | 40790-V08B |
| H1N1 | NP | Wisconsin x Victoria 2019 | Sino Biological | 40774-V08B |
| H3N2 | NP | Cambodia 2020 | Sino Biological | 40778-V08B |
| H3N2 | NP | Hong Kong 2019 | Sino Biological | 40753-V08B |
| Yamagata | NP | Phuket 2013 | Sino Biological | 40500-V08B |
| H5N1 | NP | Cambodia 2013 | Sino Biological | 40947-V08B |

**Table S2 Antigens included within Influenza multiplex binding assay.**
